# Diurnal Regulation of SOS Pathway and Sodium Excretion Underlying Salinity Tolerance of *Vigna marina*

**DOI:** 10.1101/2024.03.26.586888

**Authors:** Yusaku Noda, Fanmiao Wang, Sompong Chankaew, Hirotaka Ariga, Chiaki Muto, Yurie Iki, Haruko Ohashi, Yu Takahashi, Hiroaki Sakai, Kohtaro Iseki, Eri Ogiso-Tanaka, Nobuo Suzui, Yong-Gen Yin, Yuta Miyoshi, Kazuyuki Enomoto, Naoki Kawachi, Prakit Somta, Jun Furukawa, Norihiko Tomooka, Ken Naito

**Author notes:** **Correspondence:** K. Naito. Research Center of Genetic Resources, National Agriculture and Food Research Organization, 2-1-2 Kannondai, Tsukuba, Ibaraki, 305-8602, Japan. Authors who equally contributed to the study (Yusaku Noda, Fanmiao Wang).

## Abstract

*Vigna marina* (Barm.) Merr. is adapted to tropical marine beaches and has an outstanding tolerance to salt stress. Given there are growing demands for cultivating crops in saline soil or with saline water, it is important to understand how halophytic species are adapted to the saline environments. Here we revealed by positron emitting tracer imaging system (PETIS) that *V. marina* actively excretes sodium from the root during the light period but not in the dark period. The following whole genome sequencing accompanied with forward genetic study identified a QTL region harboring *SOS1*, encoding plasma membrane Na^+^/H^+^ antiporter, which was associated with not only salt tolerance but also ability of sodium excretion. We also found the QTL region contained a large structural rearrangement that suppressed recombination across ∼20 Mbp, fixing multiple gene loci potentially involved in salt tolerance. RNA-seq and promoter analyses revealed *SOS1* in *V. marina* was highly expressed even without salt stress and its promoter shared common *cis*-regulatory motifs with those exhibiting similar expression profile. Interestingly, the *cis-*regulatory motifs seemed installed by a transposable element (TE) insertion. Though not identified by genetic analysis, the transcriptome data also revealed *SOS2* transcription was under diurnal regulation, explaining the pattern of sodium excretion together with up-regulated expression of *SOS1*. Furthermore, we demonstrated that, under a condition of mild salt stress, the plants with the diurnally regulated SOS pathway outperformed those with the constitutively activated one.

## 1. Introduction

*Vigna marina* (Barm.) Merr. and *Vigna luteola* (Jacq.) Benth. are the most and the second most salt-tolerant species in the genus *Vigna* (Iseki et al., 2016). They are oceanic dispersal species and dominate the vegetation of tropical and subtropical beaches (Tomooka 2002). Given the increasing problems of soil salinization and freshwater shortage, it is important to understand its mechanisms of salt tolerance and the underlying genetics for developing practical salt-tolerant crops.

The preceding studies have revealed the outstanding performance of *V. marina* under salt stress. It survives 500 mM NaCl for more than 8 weeks (Yoshida et al., 2020), maintains root growth in 200 mM NaCl (Wang et al., 2023), develops thicker apoplastic barrier in response to salt stress (Wang et alk., 2024), and shows higher photosynthetic rate and stomatal conductance in 150 mM NaCl than in control (Yoshida et al., 2020). While some accessions of *V. luteola* also showed remarkable performance up to 350 mM NaCl, but others originally collected from riverbanks did not tolerate the condition of 150 mM NaCl (Chankaew et al., 2014, Yoshida et al., 2020, Wang et al., 2023). However, the sensitive accessions of *V. luteola* are still more salt-tolerant compared to *Vigna angularis* (Willd.) Ohwi & Ohashi, which is azuki bean, one of the major *Vigna* crops (Iseki et al., 2016). The following study of ours, by tracer experiment with radioactive sodium (^22^Na), revealed *V. marina* allocates the least amount of sodium in leaves, stems and roots compared to other species (Noda et al., 2022). Thus, the ability of sodium exclusion underlies its extraordinary tolerance to salt stress.

However, still unknown is whether *V. marina* simply suppresses sodium uptake from the root, or actively excretes sodium that is once loaded to xylem. Moreover, genetic factors responsible for the sodium exclusion must be identified to fully understand how *V. marina* is adapted to saline environments such as marine beaches.

Thus, in this study, we adopted the positron emitting tracer imaging system (PETIS) (Uchida et al., 2004; Fujimaki et al., 2015), which enables real-time mapping of sodium in living plants of *V. marina* and other accessions with various degrees of salt tolerance (Fig 1) (Yoshida et al., 2016) (Chankaew et al., 2014, Yoshida et al., 2020). Then we performed a forward-genetic approach by crossing *V. marina* and *V. luteola* to identify genetic loci involved in salt tolerance. We also reconstructed a new reference genome of *V. marina* and *V. luteola* and performed some comparative genomic/transcriptomic studies to identify the responsible genes and genetic variations underlying the differential expressions. The knowledge obtained in this study will provide insights regarding how to use the known genes to make a plant adapted to saline environments.

**Figure 1.**
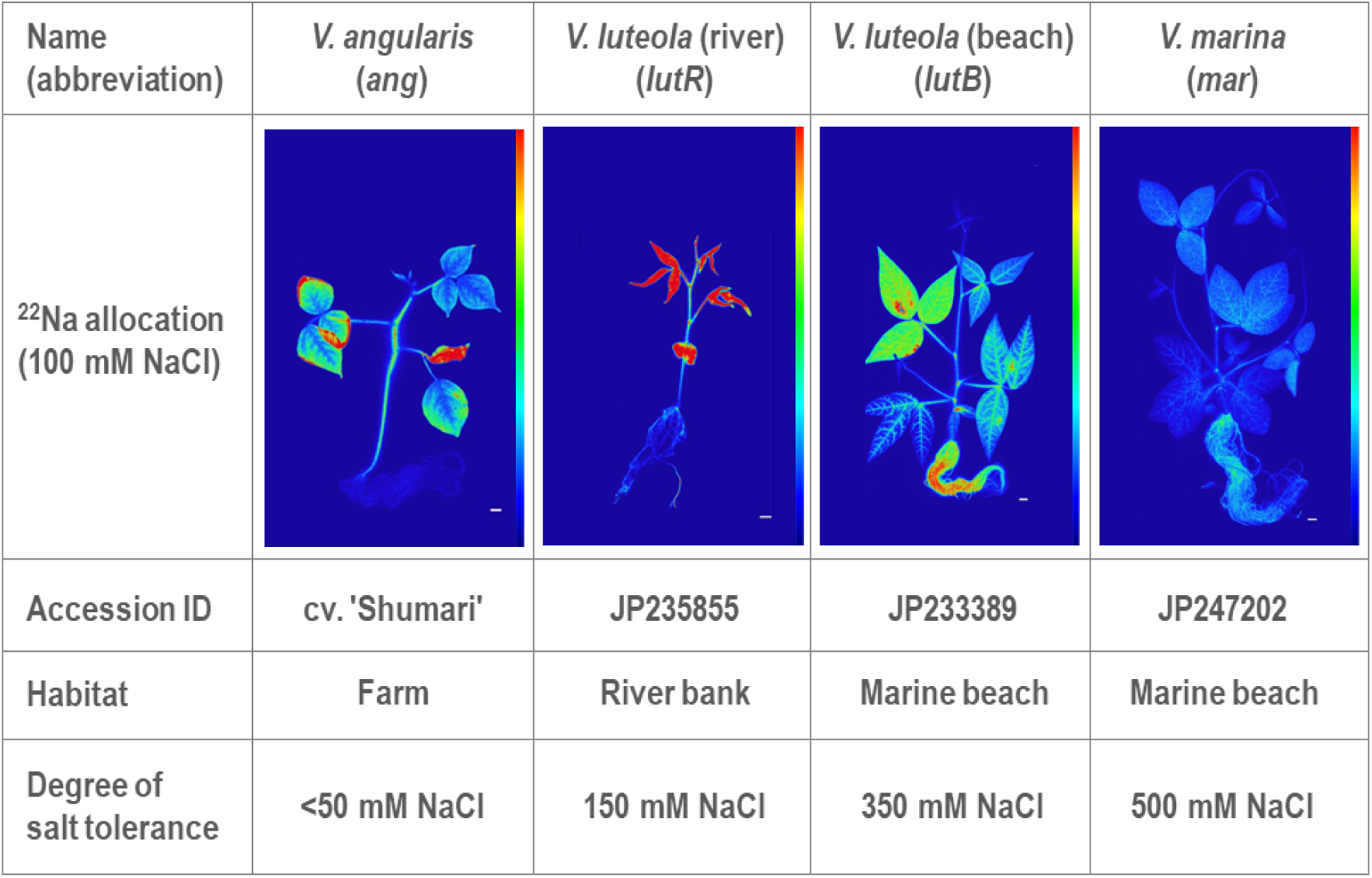
Sodium allocation and degree of salt tolerance the plant materials. Sodium allocation was visualized with autoradiography using ^22^Na. Color scales indicate relative intensity of radioactivity, where blue and red correspond to low and high, respectively.

## 2. Materials and Methods

### 2.1 Plant materials and growth condition

All plant seeds were provided by the NARO Genebank in Tsukuba, Japan (https://www.gene.affrc.go.jp/index_en.php) (Table 1). Seeds were sterilized by shaking in 70% ethanol for 5 min and then shaking another 5 min in 0.5% sodium hypochlorite. The sterilized seeds were germinated on Seramis clay (Westland Deutschland GmbH, Mogendorf, Germany) for 1 week and then transferred to hydroponic solution in a growth chamber (Light: 28°C for 14 h, Dark: 25°C for 10 h. Light intensity 500 µM-1 s-1 m-2). Each plant was cultivated for the following number of days: *V. angularis* 10 days, *V. luteola* (river) 14 days, *V. luteola* (beach) 14 days, *V. marina* 17 days. The hydroponic solution contained a diluted nutrient solution of OAT House No.1 (1.5 g L-1): OAT House No.2 (1 g L-1) (Otsuka Chemical Co, Japan) in a 1:1 ratio with concentrations of 18.6 mEq L-1 nitrogen, 5.1 mEq L-1 phosphorus, 8.6 mEq L-1 potassium, 8.2 mEq L-1 calcium and 3.0 mEq L-1 magnesium.

### 2.2 Visualization of ^22^Na allocation

To visualize ^22^Na allocation, pre-cultured plants were transferred to the Center for Research in Radiation, Isotopes and Earth System Sciences at the University of Tsukuba, Japan. The plants were transplanted into a new hydroponic solution containing 5 kBq ^22^Na (PerkinElmer, USA) with non-radioactive 100 mM ^23^NaCl. After adding the radioisotope, the plants were again incubated under long-day conditions (Light: 28°C for 14 h, Dark: 25°C for 10 h. Light intensity 200 µmol s-1 m-2) for 3 days. After incubation, the roots were carefully washed and the whole plant body was placed in a plastic bag and exposed to a storage phosphor screen (BAS-IP-MS-2025E, GE Healthcare, UK) in Amersham exposure cassettes (GE Healthcare, UK) for 24 h. The exposed screen was then scanned with a laser imaging scanner Typhoon FLA-9500 (GE Healthcare, UK). All experiments were performed independently on more than three biological replicates. To arrange radioactive intensity equally at each image, photo-stimulated luminescence and contrast were equalized by Multi Gauge version 3.0 (Fujifilm, Japan). In an experiment to visualize ^22^Na allocation in *V. marina* under different salt stresses, plants were transplanted into 100, 200 and 300 mM ^23^NaCl hydroponic solutions containing 5 kBq ^22^Na. Subsequent procedures were as described above.

### 2.3 PETIS imaging and analysis

The visualization of sodium excretion from plant roots was carried out using PETIS in Takasaki Institute for Advanced Quantum Science, Japan. Initially, a polytetrafluoroethylene sheet with holes was placed in an 8 cm high acrylic cell as a partition to enable quantification of ^22^Na radioactivity from hydroponic culture. A silicone tube was then installed for injecting the ^22^Na solution into the acrylic cell, which was attached to an acrylic board. The plants of 2 weeks old were inserted into the acrylic cell and stabilized with black urethane sponge. The plants were transferred to a growth chamber equipped with the PETIS detector. Prior to treatment with ^22^Na, the plants were pretreated with hydroponic solution containing 100 mM of non-radioactive NaCl for 1 hour. After 1 hour, the hydroponic solution was exchanged with fresh solution containing 100 mM of non-radioactive NaCl and 500 kBq ^22^Na to feed the plants for 24 hours. Then, the ^22^NaCl hydroponic solution was exchanged twice with ice-cold 10 mM CaCl_2_ solution to wash the roots. After washing, the plants were transplanted to 100 mM NaCl hydroponic solution without ^22^Na and the data collection by PETIS was started. During PETIS imaging, the surface of the hydroponic solution was maintained by a siphon pump, and roots were shaded with aluminum foil. Growth conditions during imaging were as follows: Light: 28°C for 14 h, Dark: 25°C for 10 h; light intensity: 200 μmol s-1 m-2.

PETIS collected data every minute for 48 or 72 hours. The original images acquired every minute were then re-integrated into montage images at 1 hour intervals. Quantification of ^22^Na in the hydroponic solution, root and shoot was done by defining a region of interest (ROI) with ImageJ (National Institutes of Health, Bethesda, MD, USA; http://rsb.info.nih.gov/ij/). Time-course analysis of ^22^Na radioactivity within each ROI was performed by generating a time-activity curve (TAC) from signal intensities using ImageJ.

All the experiments were independently performed more than 3 times with 3 or 4 biological replicates.

### 2.4 Genome sequencing and assembly

As described by Naito (2023), we extracted genomic DNA from the unexpanded leaves with Nucleobond HMW DNA kit (MACHEREY-NAGEL GmbH & Co. KG, Düren, Germany), size-selected the DNA with Short Read Eliminator XL kit (Pacific Biosciences of California, Inc., California,USA), and prepared 3 libraries for nanopore sequencing with SQK-LSK109 (Oxford Nanopore Technologies KK, Tokyo, Japan). Each library was then loaded to MinION R9.4.1 Flowcell (Oxford Nanopore Technologies KK, Tokyo, Japan) and was run for 72 h. The obtained raw data was transformed into fastq format with bonito-0.2.1. We also obtained short-reads with Hiseq 4000 (Illumina, San Diego, USA), which was provided as a customer service by GeneBay, Inc. (Yokohama, Japan).

For draft assembly, we used necat-0.0.1 (Chen et al., 2021) with default parameters except “PREP_OUTPUT_COVERAGE=60” and “CNS_OUTPUT_COVERAGE=40”. The contigs were polished twice with racon-1.4.3 (Vaser et al., 2017) and once with medaka-1.0.3 (https://github.com/nanoporetech/medaka).

The polished contigs were then scaffolded into pseudomolecules by anchoring to the linkage map (see below). When we found controversies between contigs and the linkage map, we manually fixed the misassemblies as described by Sakai et al. (2015). The pseudo molecules of *V. marina* were then annotated with ORFs exactly as described in our previous study (Naito et al., 2021). Those of *V. luteola* were annotated by lifting those of *V. marina* with GeMoMa-1.9 (Keilwagen et al., 2018), as described in another study of ours (Itoh, et al., 2023). We also annotated transposable elements (TEs) using EDTA-1.9.6 (Ou et al., 2019) with default parameters.

### 2.5 QTL analysis

We crossed *V. marina* and *V. luteola* (beach), selfed the F_1_ plant, and obtained F_2_ seeds. To generate replicates, the F_2_ plants were cultivated in hydroponic culture as described in Yoshida et al. (2016) for 8 weeks, and were clonally propagated by cutting (Supplementary Fig 1). For phenotyping, three clones of each F_2_ plant were cultivated in hydroponic culture with 50 mM NaCl for a week, transferred to the new culture containing 350 mM NaCl for 8 weeks, and then evaluated for 3 traits, “wilt score”, “shoot generation score” and “stay green score”. For the wilt score, the F_2_ clones were scored as 1, 3, 5, 7, or 9 according to the salt damage as shown in Supplementary Fig 1. For the shoot generation score, the clones were scored as 3 if they kept growing and generating new branches, or 1 if otherwise. For the stay green score, the clones were scored as 3 if all the leaves stayed green during the experiment, or 1 if otherwise. For genotyping, genomic DNA was extracted from young leaves of each F_2_ plant with CTAB method. The extracted DNA was then used for preparing restriction site-associated DNA sequencing (RAD-seq) (Baird et al., 2008) libraries as described by Matsumura et al. (2014), except *Bam*HI and *Mbo*I were used for DNA digestion. The prepared library was sequenced with Hiseq2000 (Illumina), which was provided as a customer service by GeneBay Inc. (Yokohama, Japan). The obtained sequence data were mapped to the *V. marina* genome with bwa (Li et al., 2010) and then processed with Stacks-1.19 (Catchen et al., 2013) to obtain genotypes. The obtained genotype data was then analyzed for linkage with onemap (Taniguti et al., 2023) to construct a linkage map, which was also used for scaffolding contigs (see above). The association between the phenotypes and the genotypes was then estimated with R/qtl2 (Broman et al., 2019).

### 2.6 Transcriptome analysis

1st experiment:

Before initiation of salt stress, plants of *V. marina* were grown in hydroponic culture for 2 weeks and pre-treated with 50 mM NaCl for 3 days. Then, at 11 am, we transferred the plants to hydroponic culture containing 200 mM NaCl. After the initiation of salt stress, we collected leaves and roots of 3 plants at 24, 36, 48 and 60 hours. The collected leaves and roots were immediately wrapped with foil and frozen in liquid nitrogen. From the collected samples, total RNA was extracted with RNeasy Plant Mini Kit (Qiagen KK, Tokyo, Japan). From the extracted RNA we prepared libraries for 3’mRNA-seq with Collibri 3’mRNA Library Preparation Kit for Illumina Systems (Thermo Fisher Scientific K.K., Tokyo, Japan). The prepared libraries were sequenced with Hiseq4000 as a customer service of GeneBay Inc. (Yokohama, Japan). After sequencing, we estimated the read counts of each gene of *V. marina* by kallisto-0.45.2 (Bray et al., 2016). With edgeR (Chen et al., 2016), we separately analyzed the leaf dataset and the root dataset. We first normalized the read counts and extracted differentially expressed genes, which were not only significant as ANOVA, but also at least one of the data points had 2-fold or greater difference compared tothe average. The extracted gene sets were then clustered with SOM-clustering (Wehrens et al., 2007).

2nd experiment:

*V. angularis, V. luteola* (river), *V. luteola* (beach) and *V. marina* were grown, salt-stressed, sampled and sequenced as described above. The RNA-seq data was quantified using the quasi-mapping-based mode of Salmon (Patro *et al*., 2017) with indexing of a de-novo assembly primary transcript of *V. marina*. The resulting raw counts were TMM (Trimmed mean of M values)-normalized using EdgeR (Robinson *et al*., 2010) and the resulting TMM-normalized values were further clustered using SOM (Wehrens *et al*., 2007) with a map size of 8 × 8.

### 2.7 Promoter analysis

All the promoter analysis was done with MEME Suite (Baily et al., 2015). First, we extracted the insertion sequence found in the upstream of *VmSOS1* and scanned it with meme command (Bailey et al., 1994) with an option of “-m mod anr”. The output was screened for Arabidopsis DAP motifs (O’Malley et al., 2016) using tomtom command with default settings.

For motif scanning, we used fimo command (Grant et al., 2011) to screen the promoters of the selected gene sets for the Arabidopsis DAP motifs and counted the numbers of “ERF1” and “CRF10” from the output. We used binominal distribution test to compare the probability between the selected gene sets vs all genes as control.

### 2.8 Transformation of *Arabidopsis thaliana*

To generate the *sos1sos2* double knockout mutant, we crossed *sos1-1* and *sos2-2* and screened the double mutant from F_2_ generation. Coding sequences of *AtSOS1* and *AtSOS2* were amplified by PCR from RAFL clone pda05112 and pda08516, a full-length cDNA clone distributed by RIKEN BRC (Seki et al., 2002), respectively. The amplified *AtSOS1* sequence was ligated downstream of CaMV 35S promoter of the binary vector pRI201-AN (Takara, Japan), and then *AtSOS2* sequences jointed with 35S promoter or 2.4 kbp upstream region of *AtPRR9* (At2g46790) were introduced in tandem with *AtSOS1* expression module (Supplementary Fig 2). These constructs were introduced into *Rhizobium radiobactor* strain GV3101. Agrobacteria were then used for transformation of *sos1sos2* plants by the floral-dip method.

### 2.9 Salinity tolerance test on Arabidopsis thaliana

We first planted seeds of Arabidopsis ecotype Col-0, *sos1/sos2* double mutant, *35S:SOS1/proPRR9:SOS2* and *35S:SOS1/35S:SOS2* transgenic plant on horticultural soil (Tachikawa Heiwa Nouen Co.Ltd, Japan) and treated them with vernalization for 3 days at 4°C. The plants were then transferred to a growth chamber set at 24°C, light 10 h, dark 14 h, light intensity 150 µM-1 s-1 m-2 and were incubated for 14 days. After that, the plants were transplanted to hydroponic culture containing 1/2 Murashige & Skoog (MS) solution and were incubated for 21 days. For salt stress, plants were pretreated with hydroponic culture with 5 mM NaCl for 3 days, and then treated with 25 mM NaCl for 1 week. After treatment, the shoots and roots were collected separately, dried at 60°C for 3 days and measured for dry weights.

## 3. Results

### 3.1 ^22^Na-allocation in *V. marina* and its relatives

In addition to our previous study that revealed the ^22^Na allocation in *V. marina, V. luteola* (beach) and *V. angularis* (Noda et al., 2022), we visualized that in *V. luteola* (river) in the condition of 100 mM NaCl (Fig 1). The result of *V. luteola* (river) showed a typical pattern of glycophytes, which was similar to *V. angularis*, presenting high ^22^Na allocation to leaves and low allocation to roots. In contrast, *V. luteola* (beach) allocated more ^22^Na to roots and less to leaves and stems, while *V. marina* highly restricted ^22^Na allocation even in roots.

### 3.2 Sodium excretion and culture alkalization by *V. marina*

To elucidate to which intensity of salt stress *V. marina* is able to keep sodium out of the plant, we fed the plants with ^22^Na in conditions of 100, 200 and 300 mM NaCl (Fig 2A). The results revealed that *V. marina* kept sodium allocation low even under 300 mM NaCl condition. Instead, we found clear traces of ^22^Na around the root of the plants in 200 mM NaCl, and more in 300 mM NaCl.

**Figure 2.**
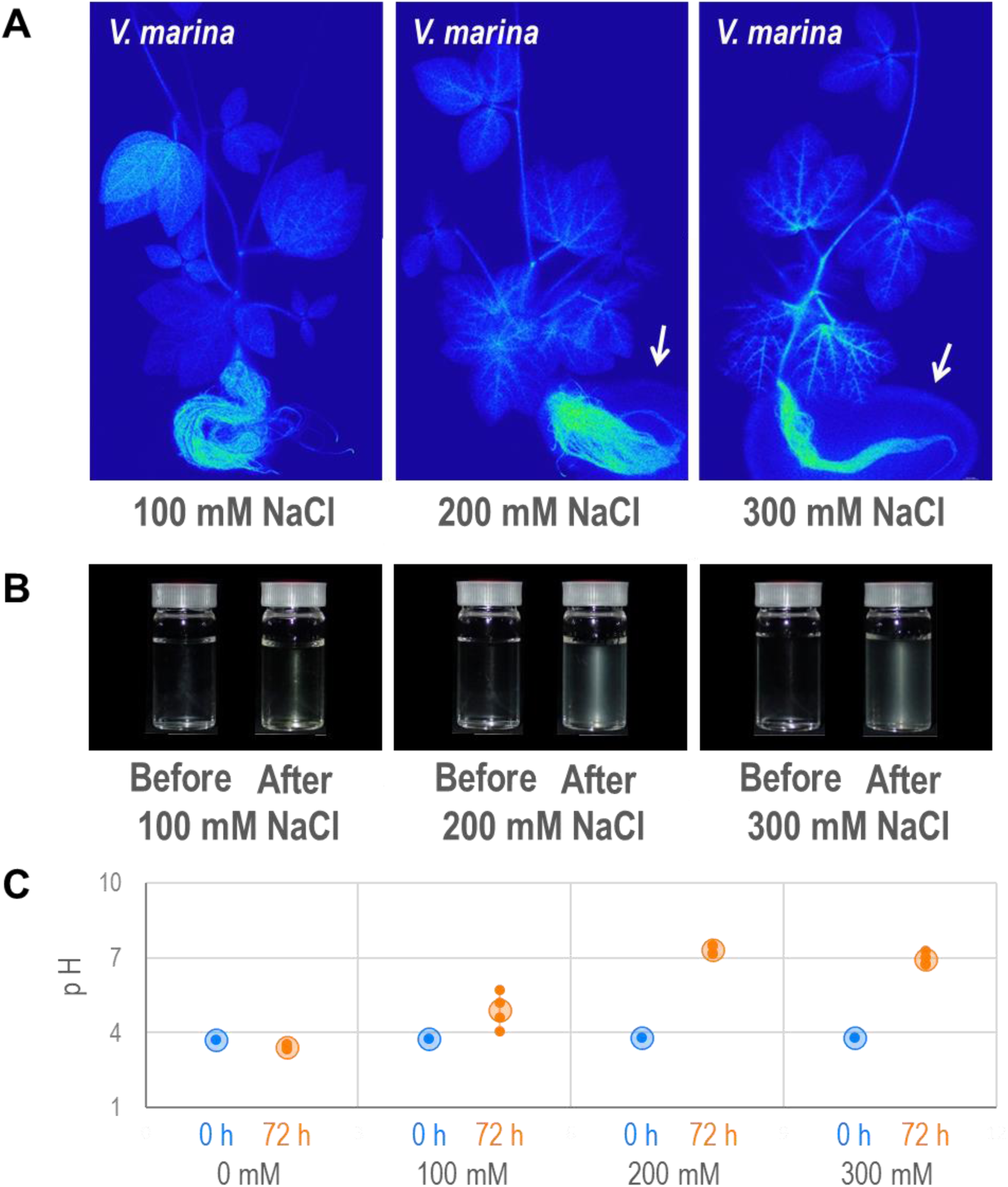
Sodium excretion and culture alkalization by *V. marina.* A. Autoradiographs of *V. marina* plants fed with 100, 200 or 300 mM NaCl containing ^22^Na. White arrows indicate traces of exudated ^22^Na from the root. B. Hydroponic culture solution before and after cultivation of *V. marina*. C. pH of hydroponic culture solution before (0h) and after (72h) cultivation of *V. marina*. Small dots indicate values of biological replicates, while larger plots and error bars indicate mean ± S.D., respectively.

During the experiment, we noticed that the hydroponic culture containing NaCl turned cloudy after plant cultivation for 72 h (Fig 2B, Supplementary Fig 3). The culture with 100 mM NaCl formed a little precipitation, while that of 200 or 300 mM NaCl formed more precipitation.

As we suspected that the precipitation was the nutrient salts because the activity of Na^+^/H^+^ antiporters could alkalize the hydroponic culture by depriving proton, we measured the pH of the hydroponic culture before and after cultivating the plants of *V. marina* (Fig 2C). As a result, with 0 mM NaCl, the plant cultivation declined the culture pH from 3.7 to 3.4, while it increased to 4.9 in 100 mM NaCl, and 7.3 in 200 mM NaCl and 6.9 in 300 mM NaCl.

### 3.3 Dynamics of ^22^Na in a living plant body of *V. marina*

To observe sodium dynamics in a plant of *V. marina*, we performed real-time mapping of ^22^Na using PETIS (Fig. 3A, Supplementary Fig 4, Supplementary Data). The real-time images revealed that, with time, the ^22^Na decreased in the root while increased in the hydroponic culture. From the images, we did not notice that the amount of ^22^Na in the shoot changed during the experiment.

**Figure 3.**
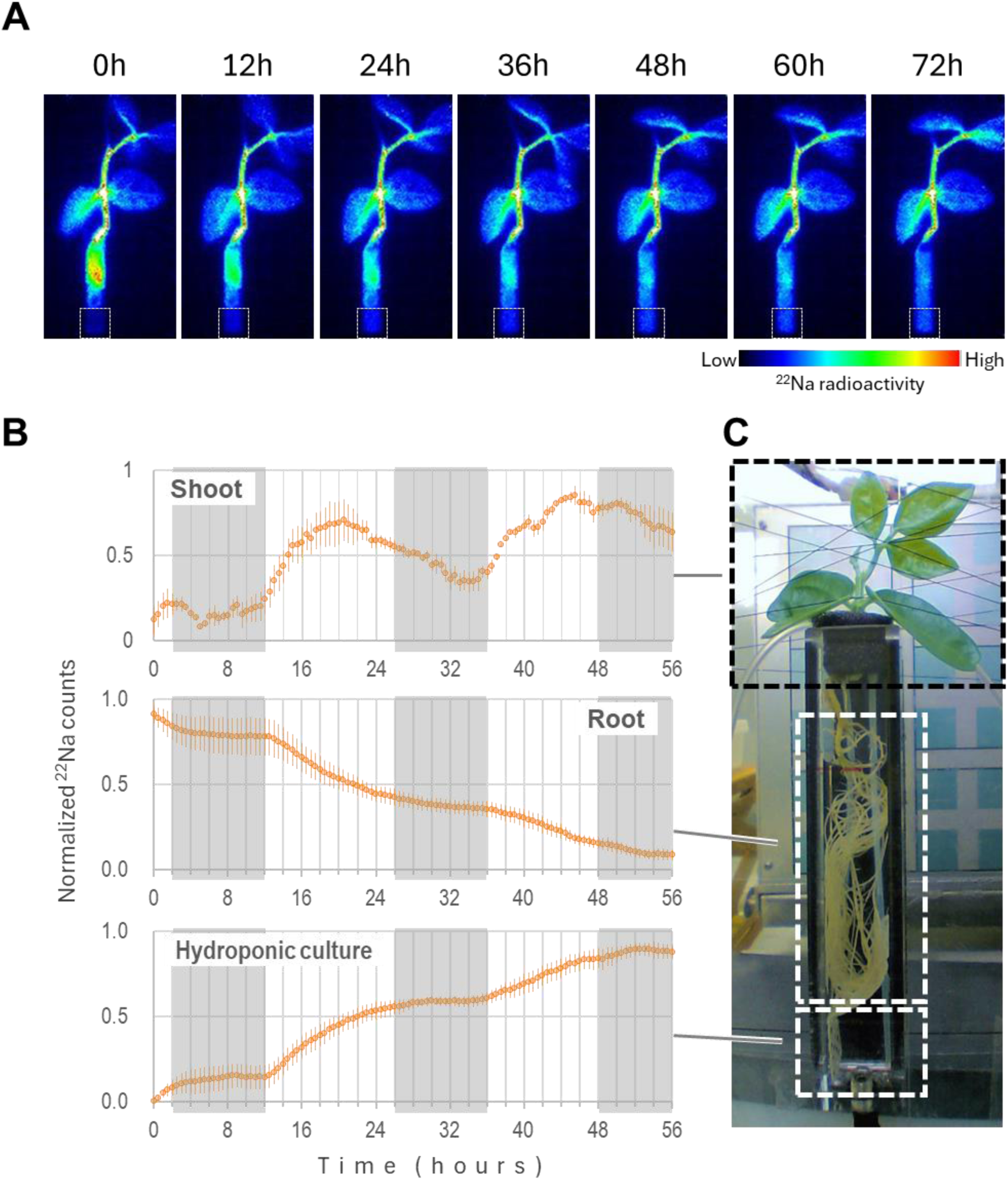
Dynamics of sodium transport in *V. marina*. A. Time-lapse images of gamma irradiation mapped by PETIS. A plant of *V. marina* was fed with ^22^Na for 24h and then time-lapsed. White boxes indicate the area of hydroponic culture measured for ^22^Na. B. Indexed amount of irradiation across time in the shoot, the root and the hydroponic culture media. Light and dark periods are indicated as white and gray in the plot, respectively. Error bars indicate standard errors. C. Setup of PETIS.

However, the following quantitative analysis revealed there was a diurnal rhythm in sodium dynamics in *V. marina.* In the shoot, the amount of ^22^Na increased during the light period and decreased during the dark period (Fig 3B), while that in the root kept simply decreasing but faster in the light period and slower in the dark period (Fig 3B). In contrast to the root, the amount of ^22^Na in the hydroponic culture increased faster during the light period and slower or even stopped during the dark period (Fig 3B). Notably, the rate of ^22^Na increase in the shoot and in the hydroponic culture was maximized in the first 6 h of the light period, indicating *V. marina* actively excreted sodium against the direction of water uptake.

### 3.4 Whole genome sequence of *V. marina* and *V. luteola* (beach)

To facilitate genetic and genomic analyses in this study, we sequenced and assembled the whole genomes of *V. marina* and *V. luteola* (beach) using Oxford Nanopore sequencer (Supplementary Table 1, Supplementary Fig 5A). The draft assembly of *V. marina* comprised 551.5 Mbp in 72 contigs with N50 length of 20.7 Mbp, whereas that of *V. luteola* (beach) was 515.7 Mbp in 146 contigs with N50 length of 18.8 Mbp. We also crossed the two accessions, obtained 286 F_2_ plants that were genotyped by RAD-seq (Supplementary Tables 2,3). With the 286 marker loci obtained, we constructed a high-density genetic map with 11 linkage groups, which corresponded to the karyotype of most Vigna species (2n = 22) (Supplementary Fig 5B). We then anchored the draft assemblies to the genetic map and reconstructed the 11 pseudomolecules with high-quality gene annotation covering more than 98% of BUSCO genes in the Fabaceae family (Supplementary Fig 1A). The genomes of the two accessions were almost colinear to each other, except inversions on chr1, chr4 and chr7 (Supplementary Fig 5C).

### 3.5 Diurnally-regulated genes in *V. marina*

As *V. marina* alkalized the hydroponic culture when exposed to salt stress, we suspected that the SOS-related genes were involved in sodium excretion. The diurnal regulation of sodium excretion as observed above further intrigued us to test whether the SOS-related genes were also diurnally regulated or not. Thus, we performed RNA-seq on the leaf and the root samples collected in the light period (11 am) and in the dark period (11 pm). The following clustering identified 4 clusters from each dataset, which were diurnally up-regulated or down-regulated at least under the condition of salt stress (Supplementary Fig 6).

Although the output of GO-enrichment analysis was not highly informative (Supplementary Tables 4-11), we found that *VmSOS2* was, both in the leaf and the root, one of the diurnally-regulated genes that were up-regulated in the light period and down-regulated in the dark period (Supplementary Fig 6). In addition, those up-regulated in the leaf during the dark period were enriched with GOs related to vesicle transport, which could be related to phloem transport (Supplementary Table 5).

### 3.6 Comparing dynamics of ^22^Na excretion among the 4 accessions

The diurnal regulation of sodium excretion of *V. marina* led us to wonder if it was a unique feature of *V. marina*. Thus, we further performed PETIS experiments on *V. angularis*, *V. luteola* (river) and *V. luteola* (beach) along with *V. marina*. As in the former PETIS experiment, we fed the plants with ^22^Na for 24 h and then transferred to non-^22^Na containing culture (Fig 4, Supplementary Figs 7-11, Supplementary Data).

**Figure 4.**
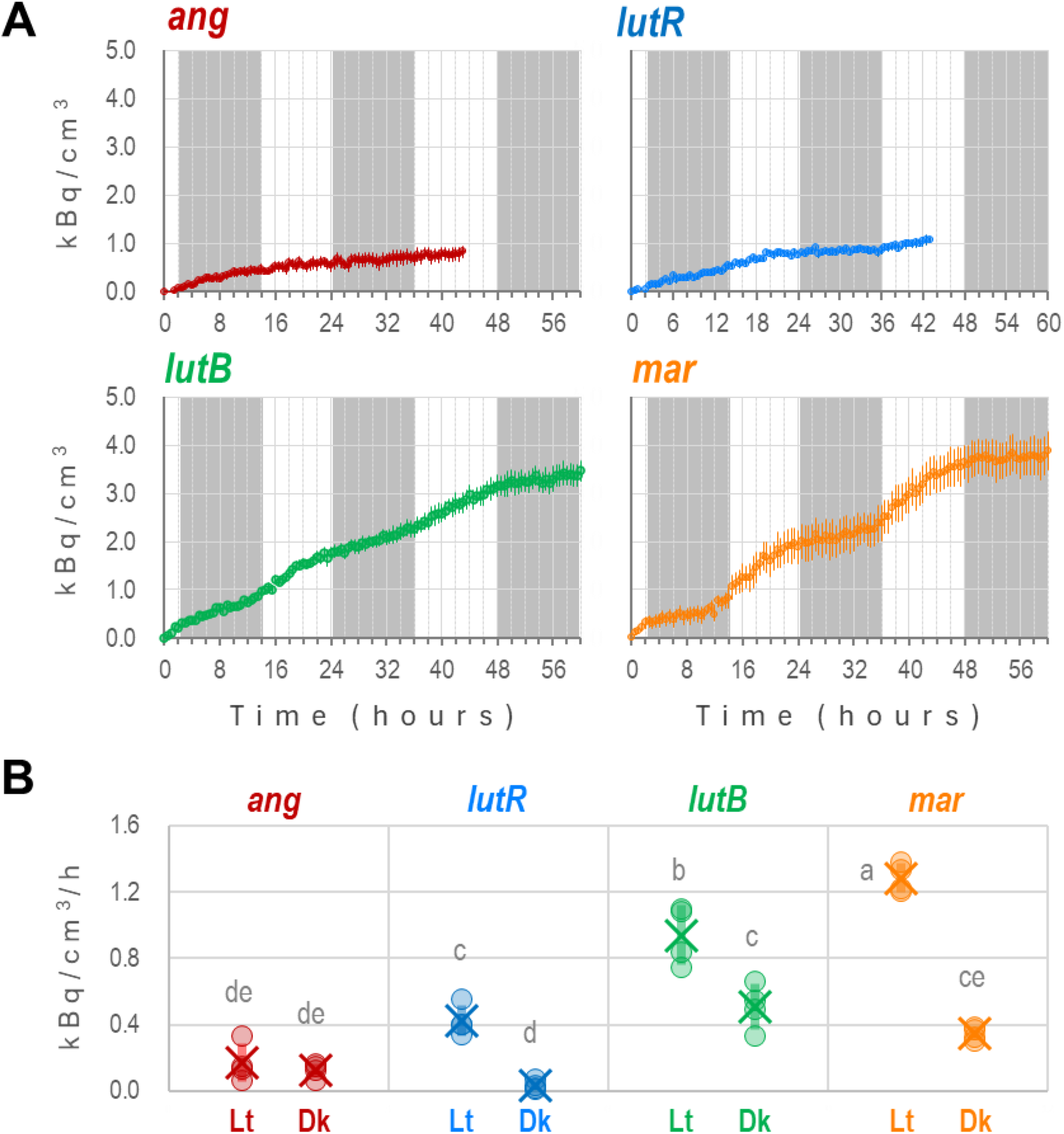
Dynamics of sodium excretion by the 4 accessions. A. Concentration of ^22^Na in the culture media across time. X- and Y-axes indicate time and the estimated concentration of ^22^Na in the hydroponic culture. Light and dark periods are indicated as white and gray in the plot, respectively. Error bars indicate standard errors. B. Velocity of ^22^Na excretion in the light period (Lt) and the dark period (Dk). The light and the dark period correspond to 12-26h and 26-36h, respectively. Different alphabets above the plots indicate significant difference by Tukey-Kramer test (p<0.05).

The real-time images by PETIS revealed a great variation across the 4 accessions (Fig 4, Supplementary Figs 8-11). *V. angularis* and *V. luteola* (river) retained little ^22^Na in the root, indicating most of ^22^Na had passed through the root and was transported to the shoot (see also Supplementary Figs 8, 9). In contrast, *V. luteola* (beach) allocated the highest amount of ^22^Na in the root. *V. marina* also allocated a high amount of ^22^Na but lower than *V. luteola* (beach) did. With time, we observed in all the 4 accessions that the amount of ^22^Na in the root decreased while that in hydroponic culture increased (Supplementary Figs 8-11, Supplementary Data).

The following quantification of PETIS data revealed that, interestingly, the diurnal pattern of ^22^Na excretion was observed not only in *V. marina*, but in *V. luteola* (beach) and even in *V. luteola* (river). (Fig 4). In contrast, *V. angularis* did not show any sign of the diurnal pattern (Fig 4). At least from 16-30 h in the PETIS analyses, the velocity of sodium excretion during the light period was significantly larger than the dark period in *V. marina* and *V. luteola*s (Fig 4B). The excretion velocities (kBq·cm^−3^·h^−1^) in the light and the dark periods were 1.3 vs 0.35 in *V. marina*, 0.94 vs 0.50 in *V. luteola* (beach), 0.42 vs 0.03 in *V. luteola* (beach), and 0.17 vs 0.12 in *V. angularis*, respectively. In addition, the velocities during the light period were significantly different across the accessions.

However, there were still differences between the accessions of *V. marina* and *V. luteola*. First, *V. luteola* (beach) and *V. marina* excreted more ^22^Na than *V. luteola* (river) did (Fig 4). Second, the difference between the light and dark periods was greater in *V. marina* than that in *V. luteola* (beach) (Fig 4B). That is, the net amount of sodium excretions was the highest in *V. marina* during the light period, but not during the dark period.

### 3.7 QTL analysis on salt tolerance

To elucidate the genetic factors underlying the salt tolerance of *V. marina*, we also evaluated the degree of salt tolerance of the F_2_ plants (wilt score, shoot generation score and stay green score) (Supplementary Fig 1, Supplementary Table 12). Together with the genotype data described above (Supplementary Tables 2,3), we performed a genome scan and detected 2 QTL peaks that were associated with all the three phenotype scores, explaining ∼25% of phenotypic variance (Fig 5, Supplementary Fig 12). In both QTLs, the alleles of *V. marina* type were associated with higher salt tolerance.

**Figure 5.**
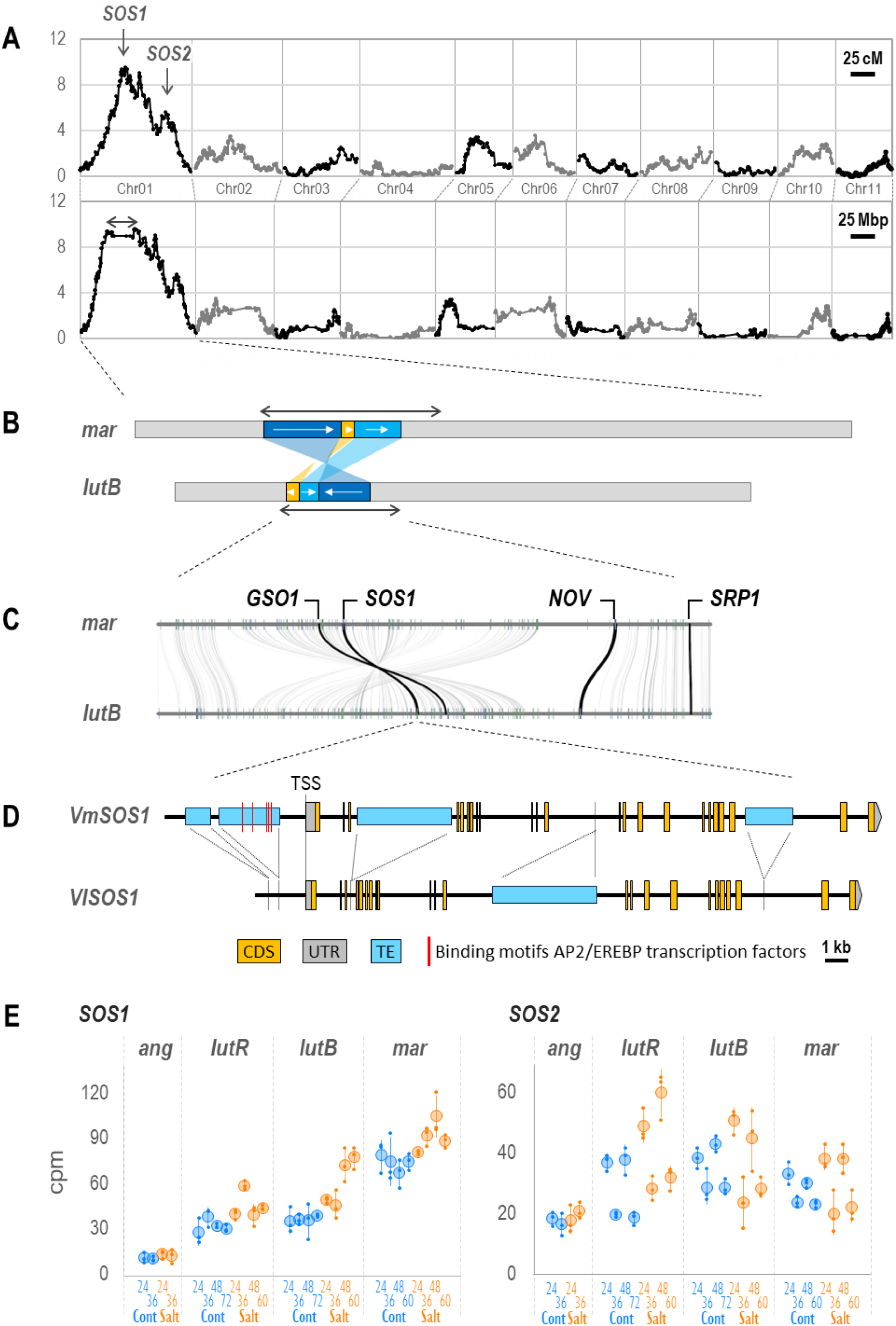
Genetic factors underlying difference of salt tolerance between *V. marina* and *V. luteola* (beach). A. QTL plot representing LOD scores. The X-axes indicate genetic distance (top) or physical distance (bottom). The two-end arrow indicates a genomic region with suppressed recombination. B. Structural rearrangement within the QTL region. Two-end arrows indicate the regions corresponding to the one in A. The arrows in the rearranged blocks indicate the corresponding order between the parents. C. Gene-based synteny plot in the rearranged region. Curved lines indicate orthologous pairs while vertical bars indicate ORFs in forward (blue) and reverse (green) directions. D. Schematic of *VmSOS1* and *VlSOS1*. Dashed lines indicate the corresponding locations between the two. E. Expression profiles of *SOS1* and *SOS2* in the root of the 4 accessions. Numbers below the X-axes indicate hours after initiation of salt stress. Cont (blue) and Salt (orange) indicate the control and salt-stressed conditions, respectively. Y-axes indicate the abundance of transcripts in count per million (CPM). Small plots indicate values of each replicate while larger ones and error bars indicate mean ± S.D.

Although the LOD curves along the genetic map presented sharp peaks, those along the physical map revealed that especially the higher peak (LOD score of) ranged at least 14.5 Mbps (17.5-32.0 Mbp). Interestingly, this region was overlapped with one of the inversions between the genomes of *V. marina* and *V. luteola* (beach) (Fig 5B, Supplementary Fig 3C) and this was why recombination was highly suppressed (Fig 5A, Supplementary Table 3). This QTL region contained several loci potentially involved in salt tolerance, including *Salt Overly Sensitive 1* (*SOS1*), *Altered Expression of Ascorbate Peroxidase 2 8* (*ALX8*) and *GASSHO1* (*GSO1*), *No Vein* (*NOV*), and *Small Rubber Particle Protein 1* (*SRP1*) (Fig 5C, Supplementary Tables 14, 15).

The other QTL region ranged ∼4 Mbp (64.3-67.6 Mbp) harboring *SOS2, Maintenance of Photosystem II Under High Light 2* (*MPH2*), *HVS22 Homologue A* (*HVA22A*), and *Purple Acid Phosphatase 26* (*PAP26*) (Supplementary Tables 13, 15).

As *V. marina* seemed to have higher Na^+^/H^+^ antiporter activity, we further investigated the polymorphisms of *SOS1* locus, which encodes one of the most well-known Na^+^/H^+^ antiporters, we further investigated the polymorphisms of this locus between the parental accessions (Fig. 5D). As a result, we found that 4 insertions of transposable element (TE)-like sequences in *VmSOS1*, whereas 1 in *VlSOS1.* Of the 5 insertions, 2 were located in the upstream of the transcription start site (TSS) of *VmSOS1*.

### 3.8 Expression profiles of the genes in QTLs

To identify differentially expressed genes (DEGs) within the QTL regions, we performed another transcriptome analysis on the 4 accessions. In this analysis, we started salt stress at 11:00 am on day0 and sampled 11:00 am (24h) and 11:00 pm (36h) on day1, and 11:00 am (48h) and 11:00 pm (60h) on day2. For *V. angularis,* we sampled only on day1 (24 and 36h).

Of all the genes in *V. marina*, we first looked into the expression profiles of *SOS1* and *SOS2*, as the results described above (Figs 2-4, 5A-D) indicated *SOS1* and *SOS2* were involved in Na^+^ excretion and its diurnal regulation. As expected, *SOS1* showed a positive correlation between the salt tolerance and the transcript abundance in the root (Fig. 5E, Supplementary Tables 12-15). In *V. angularis,* the transcript of *SOS1* was the least in the control condition and was not induced by salt stress. In *V. luteola* (river), it was ∼3 times higher than that of *V. angularis* in the control condition and was slightly increased in response to salt stress. In *V. luteola* (beach), it was similar to that of *V. luteola* (river) in the control condition, but more strongly induced in response to salt stress. In *V. marina*, it was already high in the control condition and even increased in response to salt stress (Fig 5E). The expression pattern of *SOS1* in the leaf was similar to that in the root, except it was dramatically induced in *V. luteola* (river) in response to salt stress (Supplementary Table 14). Further, the independent qPCR analysis confirmed the expression profile of *SOS1* (Supplementary Fig 13). *SOS2* was again up-regulated in the light period and down-regulated in the dark period, not only in *V. marina* but also in *V. luteola* (river) and *V. luteola* (beach) (Fig. 5E, Supplementary Tables 12-15). In contrast, there was no sign of diurnal regulation in *V. angularis* (Fig. 5E).

As those in the QTL regions other than *SOS1* and *SOS2* could be responsible for the salt tolerance of *V. marina*, we performed SOM-clustering and grouped them according to the expression patterns. As a result, *ALX8*, *NOV*, *SRP1*, *HVA22A*, *MPH2* and *PAP26* were grouped into a cluster containing *SOS1*, which were more abundantly transcribed in the root of *V. marina* than in *V. luteola*s under any conditions and timepoints (Supplementary Tables 13, 15). Of them, *SOS1, ALX8, HVA22, MPH2* and *PAP26* were also highly transcribed in the leaf of *V. marina* before salt stress, although some in *V. luteola* (river) showed a dramatic increase by salt stress (Supplementary Tables 12, 14). Though *GSO1* was not transcribed much in the control condition, it was strongly up-regulated in the root by salt stress (Supplementary Tables 13, 15).

### 3.9 Promoter analysis on *SOS1* and *SOS2*

Because we considered the higher expression of *SOS1* is one of the key factors for the salt tolerance of *V. marina*, we investigated the promoter (2.5 kb upstream from TSS) of *SOS1* locus for enriched motifs.

First, as the *VmSOS1* had a TE insertion at the site of −1,267 bp from TSS, we screened the inserted sequence for enriched motifs. The results revealed “CGCCRGGCGGMCTG” as enriched, which could potentially be a binding site for AP2/EREBP transcription factors such as ERF1 and CRF10 (Figs. 5D, 6B, C).

**Figure 6.**
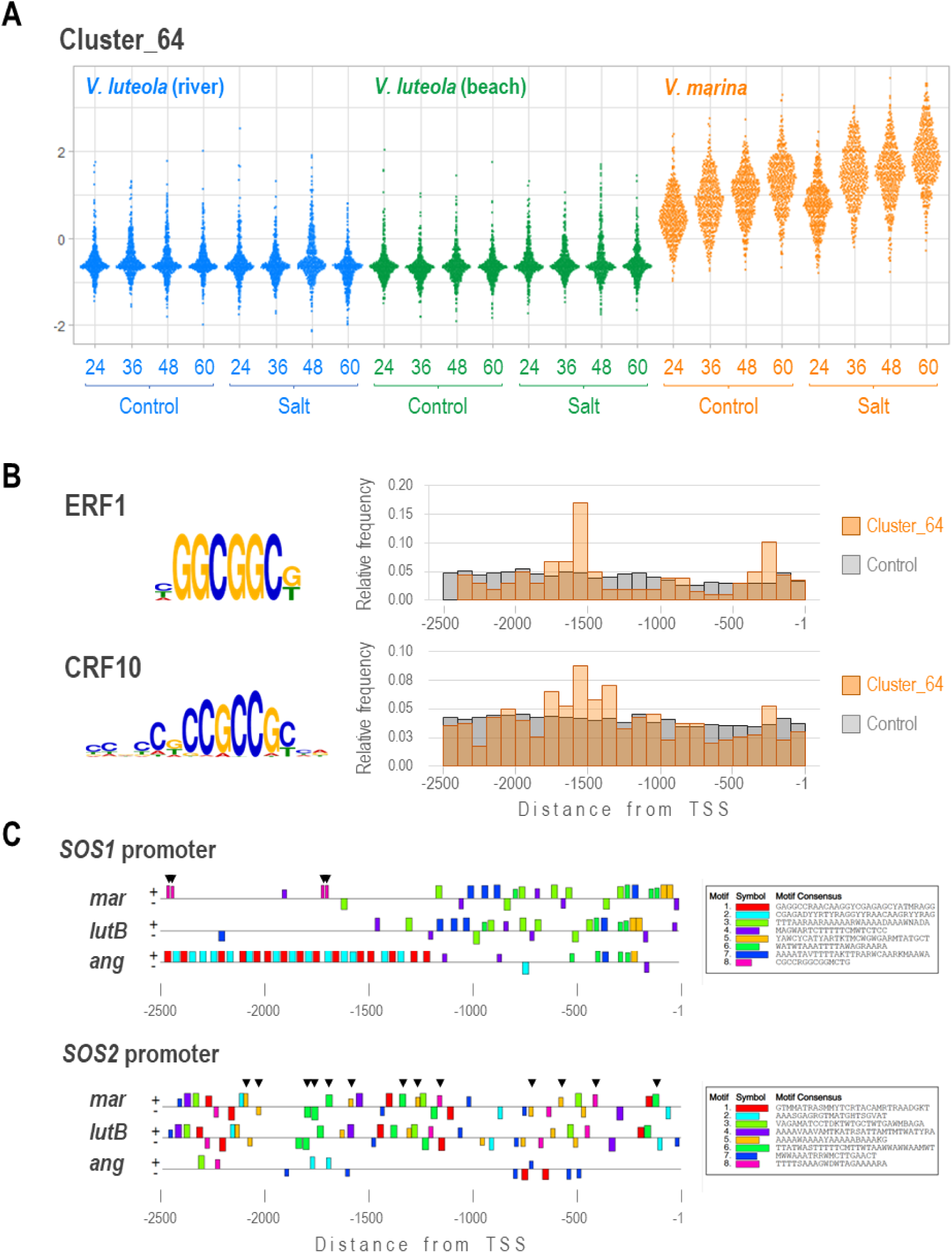
Enriched motifs in promoters (2.5 kb upstream) of *SOS1*, *SOS2* and highly expressed genes in *V. marina*. A. Expression profiles of genes in cluster_64 grouped by SOM clustering. B. Enriched motifs in the promoter sequences of the genes in cluster_64. The potentially binding transcription factors and the relative frequency of the motifs across the promoters are also shown. C. Enriched motifs of the promoters of *SOS1* and *SOS2* loci in *V. marina*, *V. luteola* (beach) and *V. angularis.* Arrowheads indicate the motifs containing AP2/EREBP binding sites in *SOS1* promoters and those potentially involved in circadian rhythm in *SOS2* promoters.

To test if this motif is associated with the expression profile of *VmSOS1*, we extracted the promoter sequences from those genes highly expressed in *V. marina* but not in *V. luteola*s (cluster_64) and from other genes as control (Fig 6A). Compared to the control genes, the promoters of genes in cluster_64 were significantly more enriched with the potential binding sites for ERF1 (0.48/2.5 kb vs 0.30/2.5 kb) and CRF10 (0.94/2.5 kb vs 0.82/2.5 kb). In addition, the ERF1 motifs were highly enriched around 0.2-0.3 kb and 1.5-1.7 kb upstream of those in cluster_64 compared, whereas it showed more flat distribution in the promoters of the control genes (Fig 6B). The CRF10 motifs were also enriched in 1.3-1.7 kb upstream of the cluster_64 genes (Fig 6B).

Finally, we compared the promoter sequences of *SOS1* and *SOS2* across *V. marina*, *V. luteola* (beach) and *V. angularis*. The motif scan revealed that the *V. angularis* conserved very few motifs, except the first ∼400 bp upstream of the *SOS1* TSS (Fig 6C), whereas *V. marina* and *V. luteola* (beach) highly conserved the promoter structures except the TE insertion (Figs 5D, 6C). Of the four “CGCCRGGCGGMCTG” motifs in the *VmSOS1* promoter, two were placed at 1,689 bp and 1,711 bp upstream of TSS, as in many of the cluster_64 genes (Figs 5D, 6B, C). The result also revealed that the promoters of *VmSOS2* and *VlSOS2* harbored 13 potential binding sites for *B-box domain protein 31* (*BBX31*), which is involved in “circadian rhythm”, whereas *VaSOS2* harbored no motifs of such kind (Fig 6C).

### 3.10 *SOS2* overexpression vs. *SOS2* diurnal expression

As we were surprised to find the conservation of the diurnally regulated sodium excretion in *V. marina* and *V. luteola*, we wondered if it has any adaptive aspect. To elucidate this query, we reproduced the *V. marina*’s expression profile of *SOS1* and *SOS2* in *Arabidopsis thaliana* by introducing *35S:SOS1* and *proPRR9:SOS2* into *sos1/sos2* double mutant, as the *proPRR9* promotes transcription in the light period but not in the dark period (Ito et al., 2007) (Supplementary Fig 2). To compare, we also tested Col-0, *sos1/sos2* double mutant and *35S:SOS1/35S:SOS2* double overexpression (Supplementary Fig 2). In the experiment, we cultivated the plants in hydroponic culture or plate culture with or without salt stress, measured the dry weight of shoots and roots separately, and calculated the relative dry weight (RDW).

The results from the hydroponic culture revealed a significant difference in the root across different patterns of *SOS1/SOS2* expression (Fig. 7A). While the shoot RDW was significantly lower only in the *sos1/sos2* double mutant, the root RDW was significantly higher in the *35S:SOS1/proPRR9:SOS2* line and the *35S:SOS1/35S:SOS2* double overexpression line. Notably, the *35S:SOS1/proPRR9:SOS2* line showed even higher RDW values than the *35S:SOS1/35S:SOS2* line.

**Figure 7.**
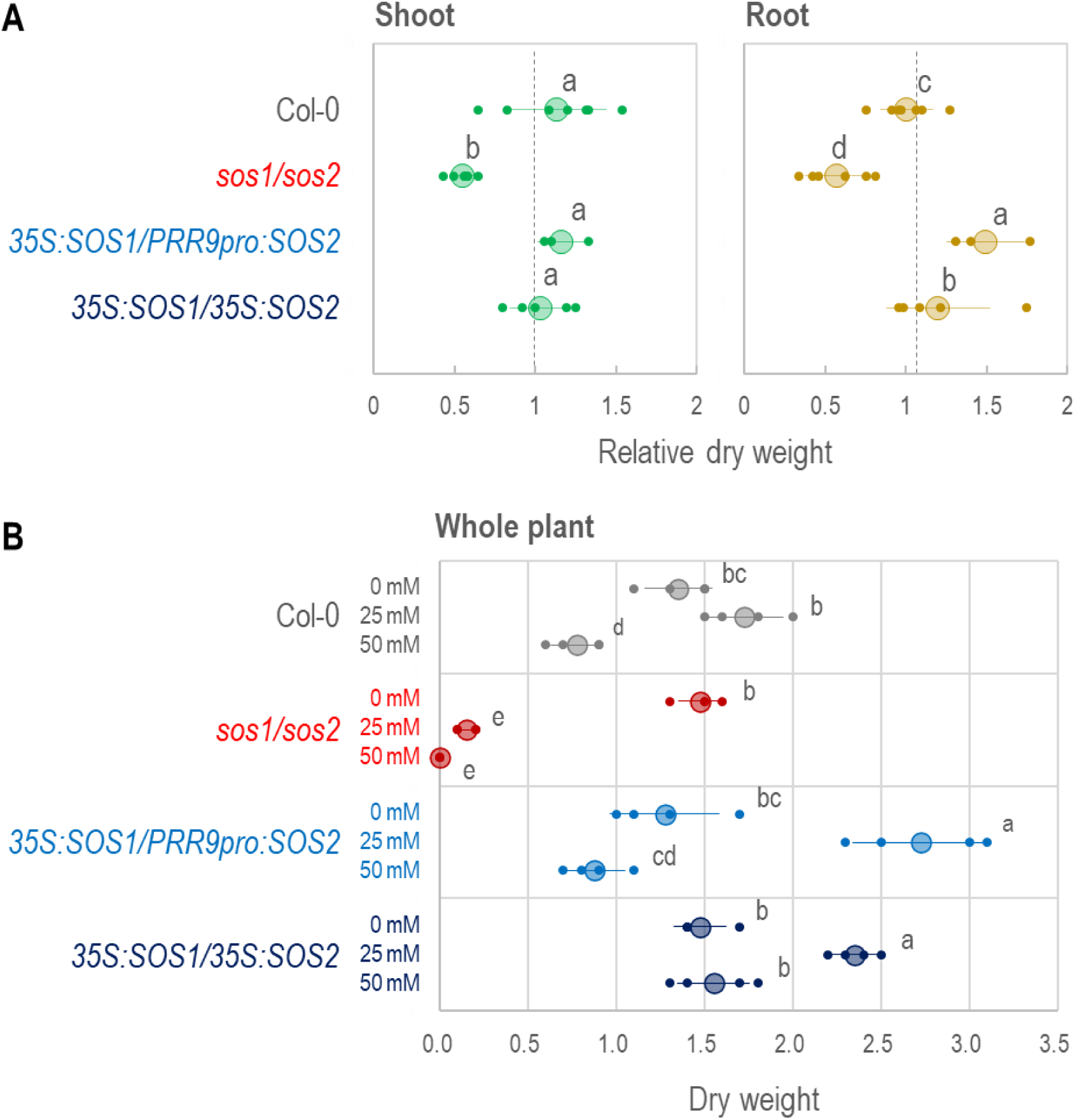
Relative dry weight of shoots and roots in various *SOS1/SOS2* expression lines. A. Relative dry weights of the shoot and the root cultivated in hydroponic culture. B. Dry weights of the whole plants cultivated on palte culture. While small plots indicate the values of each replicate, large plots and error bars indicate means ±S.D. Different alphabets above the plots indicate significant difference by Tukey-Kramer test (p<0.05).

In the plate culture, the *35S:SOS1/proPRR9:SOS2* line also outperformed Col-0 and *sos1*/*sos2* lines at least in the condition of 25 mM NaCl, though not significantly different from the *35S:SOS1/35S:SOS2* line (Fig 7B). In addition, the *35S:SOS1/35S:SOS2* line showed the best performance in the condition of 50 mM NaCl (Fig 7B).

## 4. Discussion

Here we revealed not only that *V. marina* has greater ability to excrete sodium out of the root, but that the sodium excretion is under diurnal regulation. As the sodium excretion must occur against the direction of water uptake, it was surprising to observe that *V. marina* excreted sodium mainly during the light period, which is usually the time of transpiration. In addition, the forward genetic and transcriptome analyses indicate that *SOS1* could determine the maximum rate of sodium excretion and *SOS2* could regulate its diurnal manner. The up-regulation of *SOS1* could be due to the TE insertion in the promoter, which might have installed binding motifs for AP2/EREBP transcription factors. The diurnal regulation of *SOS2* could also be explained with its promoter sequences, which had been acquired before the divergence of *V. marina* and *V. luteola.* Finally, we demonstrated that the diurnal expression of *SOS2* significantly confers salt tolerance and attenuates the negative effect of the SOS pathway on root growth, at least to some extent of salt stress.

### 4.1 Diurnal regulation of sodium excretion

It was surprising to discover by PETIS that *V. marina* and *V. luteola* excrete sodium in a diurnally-regulated manner: more in the light period and less in the dark period (Figs 3, 4). The diurnal pattern of ^22^Na in the shoot was not surprising, as sodium is usually transported to the shoot in a transpiration-dependent manner (Dinneny, 2015) and re-circulated to the root *via* phloem transport mediated by vesicle transport (Berthomieu et al., 2003). The recirculation is supported by our results that the vesicle transport genes were up-regulated in the leaf during the dark period (Supplementary Table 5). The decrease of root sodium during the light period could also be, to some extent, due to transpiration-dependent transport to the shoot. However, as the sodium concentration in hydroponic culture also increased in the light period, the decrease in the root was also due to sodium excretion. If the outward transport were a passive diffusion, it should be slow in the light period and fast in the dark period, given the transport must occur against the direction of water uptake. Given the result was the opposite (Figs 3, 4), *V. marina* is able to actively excrete sodium from the root during the light period.

### 4.2 Role *SOS1* and *SOS2* in sodium excretion of *V. marina*

As the sodium excretion was accompanied by alkalization of hydroponic culture (Fig 2), we consider the Na^+^/H^+^ antiporter activity of *SOS1* (Ismail and Horie, 2017) plays the key role in excreting sodium out of the root. This hypothesis is also supported by our results that *SOS1* is located in the highest QTL of salt tolerance (Fig 5) and that *SOS1* transcription was highly correlated not only with salt tolerance but the ability of sodium excretion (Figs 3-5). In addition, the pattern of *SOS1* expression also well explains the manner of salt allocation observed in this study (Fig 1A) and our previous study (Noda et al., 2022): The lower expression of *SOS1* in the root of *V. angularis* and *V. luteola* (river) cannot suppress sodium loading to xylem and allow high allocation to the shoot. In contrast, the higher expression of *SOS1* in the root of *V. marina*, even before salt stress, enables immediate sodium excretion after initiation of salt stress. Although the expression of *SOS1* in the root of *V. luteola* (beach) is strongly induced, the lower expression in the control condition might allow sodium to accumulate in the root in the early stage of salt stress.

The diurnally-regulated transcription of *SOS2*, encoding calcineurin binding protein-like (CBL) interacting protein kinase (CIPK) that directly phosphorylates and activates SOS1 (Liu et al., 2000), also explains the observed pattern of sodium excretion. Though we have found only correlation, but *V. marina* and *V. luteola*s exhibited a diurnal pattern in *SOS2* transcription and sodium excretion, while *V. angularis* did neither in SOS2 transcription nor sodium excretion (Figs 3-5). However, although *VmSOS2* was also located within one of the QTLs, it could not explain the difference of salt tolerance between *V. marina* and *V. luteola* (beach) given its transcriptional pattern was not different (Fig 5).

### 4.3 Evolution of transcriptional regulation in *SOS1* and *SOS2*

Our results suggested that up-regulation of *VmSOS1* is a new example of gene regulatory network rewiring by TE insertions (Feschotte 2008, 2017). The TE insertion at −1,207 bp position of *VmSOS1* placed the AP2/EREBP motif at −1,689 bp (Figs 5D, 6C), and such motifs were also enriched around −1.5 kbp positions of the genes highly expressed in *V. marina* (Fig 6). Though what we found is currently only a correlation, it is an intriguing hypothesis to test whether the TE has installed the key *cis*-regulatory element to the *SOS1* promoter.

The output of our promoter analysis also suggested that *V. marina* and *V. luteola* have acquired a set of *cis*-regulatory elements for diurnal regulation before they have diverged from the common ancestor. The structure of *VaSOS2* promoter, which was totally different from *VmSOS2* and *VlSOS2*, could also explain why *VaSOS2* is not regulated in the diurnal manner (Fig 5E). As the diurnal expression of *SOS2* is also conserved in *V. luteola* (river), it should have played important roles in adaptation, not only to high salinity but also to other environmental ques.

Although the SOS pathway in model plants is also diurnally regulated (Park et al., 2016, Soni et al., 2013, Kim et al., 2013), the results in this study are different from those in the preceding studies. First, the transcription of *SOS1* does not seem to be diurnally regulated in Vigna species (Fig 5E). In Arabidopsis, *SOS1* transcription is more in the light period and less in the dark period (Park et al., 2016), whereas in rice it is more transcribed in the dark period (Soni et al., 2013). Second, at least in *V. marina*, the activity of the SOS pathway is maximized in the early morning (Figs 3, 4), whereas it should be higher in the dark period in Arabidopsis and rice. In rice, *SOS2* is, together with *SOS1*, more transcribed in the dark period (Soni et al., 2013). In Arabidopsis, although there are few literatures reporting transcriptional regulation of *SOS2*, SOS2 protein is known to physically interact with *GIGANTEA* (*GI*), which is expressed during daytime, and to be prevented from activating SOS1 (Kim et al., 2013). Although *GI* is degraded by salt stress, it is a time-consuming process and thus negatively affects the SOS1 activity. Thus, the daytime-oriented activity of the SOS pathway could be a unique feature of salt tolerance in *V. marina*.

### 4.4 Role of chromosome rearrangement in fixing the adaptive haplotype within the QTL

Although we consider *SOS1* in the most important gene for the *V. marina*’s ability to excrete sodium, there are several other genes that could be involved in salt tolerance and highly expressed or salt-induced in *V. marina* (Fig 5C, Supplementary Tables 14,15). As *GSO1* (Pfister et al., 2014) and *NOV* (Tsugeki et al., 2009) are involved in development of Casparian strip and endodermis, respectively, they might contribute to developing the thick apoplastic barrier in the root of *V. marina* that effectively suppresses sodium loading to xylem (Wang et al., 2024). *SRP1* has dual roles in enhancing not only tissue growth but in tolerance to abiotic stresses (Kim et al., 2016). Thus, within the QTL regions, *V. marina* might have a set of adaptive alleles across multiple gene loci.

What is interesting in our findings is that the QTL region contains structural rearrangement spanning more than 15 Mbp (Fig 5B,C, Supplementary Fig 3). As this rearrangement almost completely suppressed recombination across the whole QTL region, more than 300 gene loci behaved like a single genetic locus in our mapping population (Fig 5A-C, Supplementary Tables 14, 15). Such a large rearrangement could contribute chromosome speciation, as rearranged chromosomes make the favorable allele combinations resilient to outcross (reviewed by Faira and Navarro, 2010). Without the rearrangement, the linkage of adaptive alleles would be easily broken and the progenies are likely to inherit less favorable set of alleles compared to the parental genotypes.

### 4.5 Positive and negative effect of the SOS pathway

It was first puzzling to notice that the diurnal regulation of SOS-dependent sodium excretion is conserved across species, given high and low tides do not occur in 24 h cycle, or extreme high tides by typhoons could occur at any time of a day. However, if the SOS pathway is energetically costly, the diurnal down-regulation could contribute to saving energy and to keeping growth.

Supporting our hypothesis, the Arabidopsis line transformed with *35S:SOS1/proPRR9:SOS2* showed the best performance under 25 mM NaCl condition (Fig 7). Although the double overexpression line (*35S:SOS1/35S:SOS2*) showed better performance than Col-0 and the *sos1/sos2* double mutant, it underperformed the *35S:SOS1/proPRR9:SOS2* line at least in the condition of 25 mM NaCl. Thus, though the SOS pathway is essential in salt tolerance, it could also negatively affect growth. The diurnal regulation seems to be a reasonable compromise to confer a certain level of salt tolerance with minimized loss of growth. It could be even more reasonable to activate the SOS pathway in the light period, given it is the time of the highest transpiration-dependent sodium uptake (Fig 3).

### 4.6 Limitations

Though we have demonstrated the importance of *SOS1* and *SOS2* in *V. marina* and *V. luteola*, we have not thoroughly investigated the second QTL for the genes that could explain the parental difference of salt tolerance. The possible candidates are *HVA22A*, *PAP26*, *MPH2*, and so on. *HVA22* is often strongly induced by stress and ABA, but its functional role in stress tolerance is not yet clear. *PAP26* is a member of purple acid phosphatase and enhances salt tolerance when overexpressed (Abbasi-Vineh et al., 2021). It does so by enhancing ion homeostasis *via* the SOS pathway, ROS scavenging ability, and acid phosphatase activity required for increasing available P storage. *MPH2* is required for repairing photosystem II (PSII) under fluctuating light conditions or photoinhibitory light (Liu and Last, 2017). As PSII is particularly vulnerable to abiotic stresses including salt stress (Jajoo, 2013), the higher expression of *MPH2* (Fig 5) could help acclimation of *V. marina* to saline environments. However, further studies are certainly needed to test whether these genes, or others, are responsible for *V. marina*’s salt tolerance.

In addition, although this study provides insights to the mechanisms of salt tolerance in *V. marina*, it is not easy to explain how *V. luteola* (beach) keeps excreting sodium even in the dark period (Fig 4). One possible explanation is that it could be passive diffusion along the cline of ^22^Na concentration across the root and the hydroponic culture, as *V. luteola* (beach) accumulates more sodium in the root (Figs 1, 5) (Noda et al., 2021, Wang et al., 2023). However, it is also possible that other mechanisms than the SOS pathway that enable active excretion of sodium for 24 h. In any case, further studies are needed to elucidate the query.

## 5. Conclusion

We revealed at least one aspect of the mechanism of salt tolerance in *V. marina*, which lives in marine beach and is the most salt-tolerant species in the genus. It has higher potential of sodium excretion from the root, presumably through the elevated expression of *VmSOS1.* In addition, the sodium excretion is diurnally regulated, *via* diurnal transcriptional regulation of *VmSOS2*, which could save the energy cost of the SOS pathway and maximize growth under saline environment. This knowledge will be valuable information for future crop development especially for cropping with saline water and soil.

## ACKNOWLEDGEMENTS

The corresponding author is grateful to Dr. Akira Isogai, Dr. Yasunobu Ohkawa, Dr. Naoko Nishizawa, Dr. Kazuo Shinozaki and other committee members of the PRESTO program for their valuable advice and extraordinary support, especially when he was involved in an incident in his affiliation. The Arabidopsis full-length cDNA clone used in this research was developed by the plant genome project of RIKEN Genomic Science Center.

## Funding Information

This study was financially supported by JST PRESTO Grant Number JPMJPR11B6, Moonshot R&D Program for Agriculture, Forestry and Fisheries by Cabinet Office, Government of Japan (JPJ009237), JSPS KAKENHI (JP18H02182), NARO Innovation Program, Environmental Radioactivity Research Network Center (Y-19-05) and Interdisciplinary Project on Environmental Transfer of Radionuclides (No. Y-1).

## Data availability

All the sequence data are available from the Sequence Read Archive of the NCBI website. The BioProject ID is PRJNA1080052 and BioSamples accessions SAMN40099256-99737.

## CONFLICT OF INTEREST STATEMENT

The authors have no conflicts of interest to declare that are relevant to the content of this article.

## Author contributions

KN, NS, JF and NT conceived the study.

KN, HO and HS sequenced, assembled and annotated the genomes.

SC, CM, EOT, YT, PS, YT, NT and KN developed the mapping population, performed phenotyping, genotyping, and QTL analysis.

YN, NS, YGY, YM, KE, NK and JF performed the tracer experiments and PETIS.

HO, YN, CM and KN collected the RNA samples and performed sequencing.

FW, HO and KN analyzed the transcriptome data.

FW and KN performed comparative genomic analysis and promoter analysis.

HA performed transformation.

NY and YI performed functional validation of the transformants.

KN and YN wrote the manuscript.

## Notes

### Competing Interest Statement

The authors have declared no competing interest.

### Summary of Updates

In addition to V. marina, we sequenced and assembled V. luteola genome and performed comparative analysis. As a result, we identified structural rearrangement spanning almost all the QTL region. The comparative analysis also enabled us to identify an TE insertion in the SOS1 promoter in V. marina, which is absent in V. luteola. The following promoter analyses revealed that the inserted TE has AP2/EREBP binding motifs, which turned out to be enriched in the promoters of the genes highly expressed in V. marina

