## Supplementary figures and images for "Diurnal Regulation of SOS Pathway and Sodium Excretion Underlying Salinity Tolerance of *Vigna marina*"

### Supplemental videos

#### Slide 1
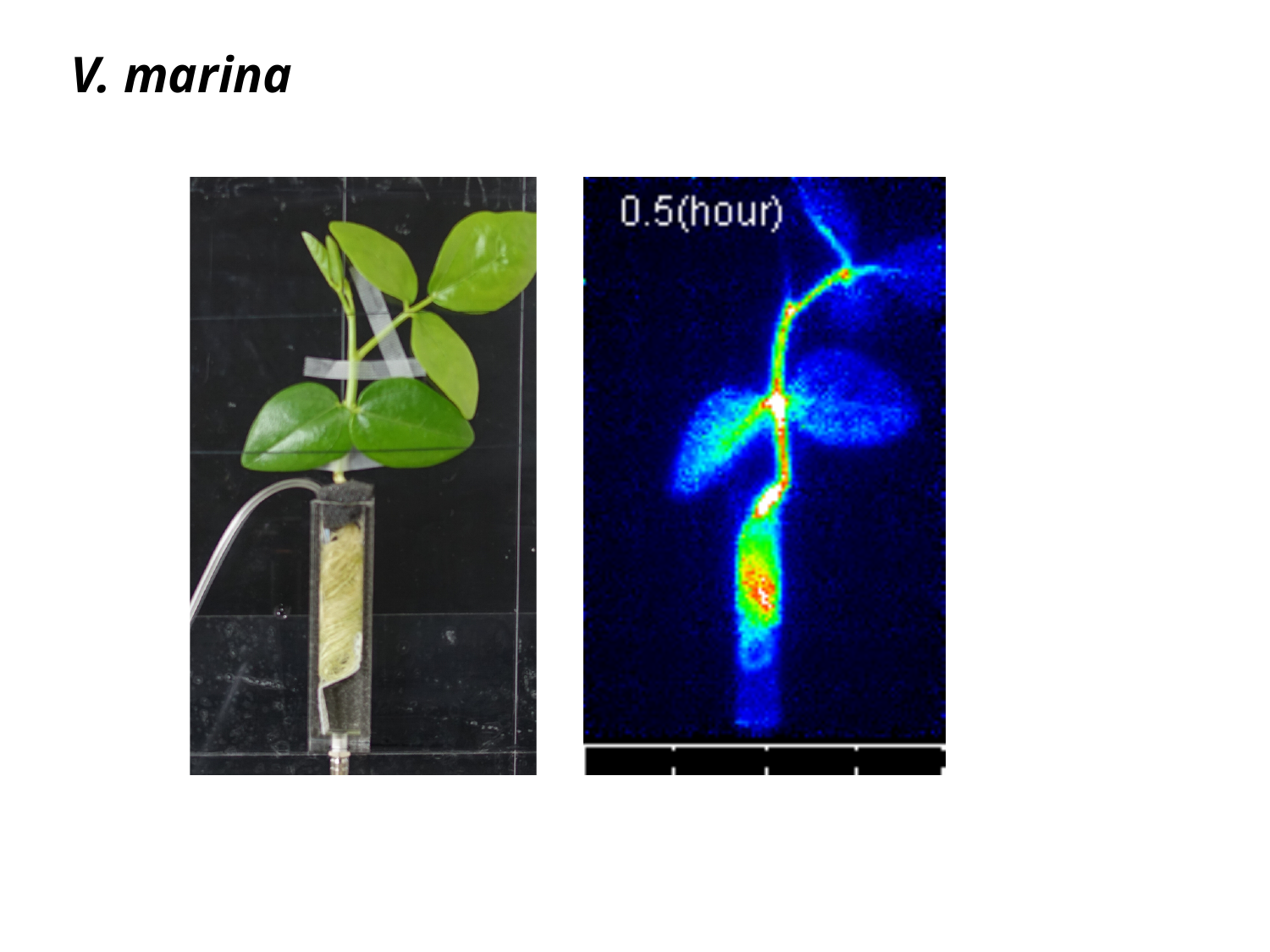

V. marina

#### Slide 2
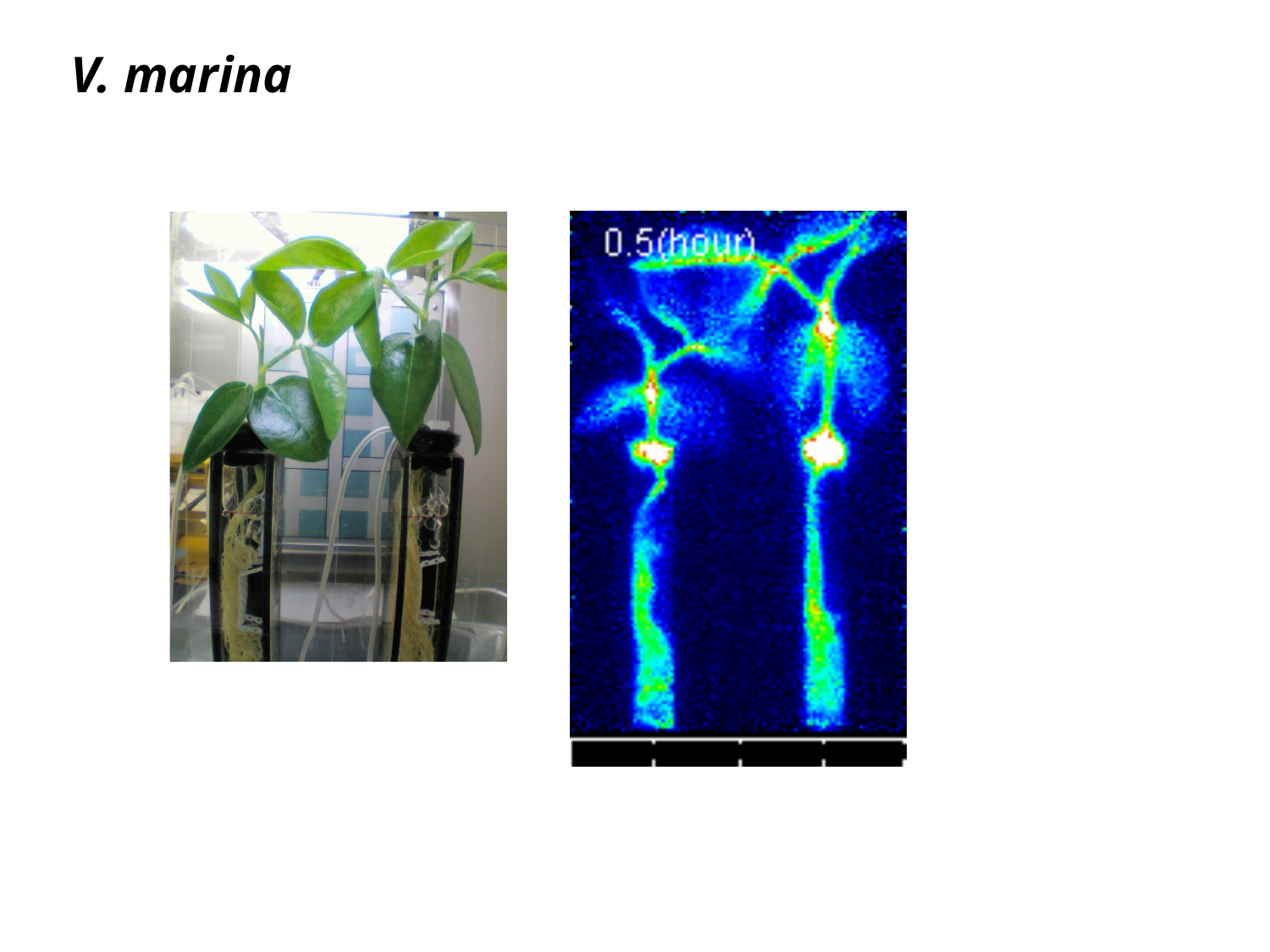

V. marina

#### Slide 3
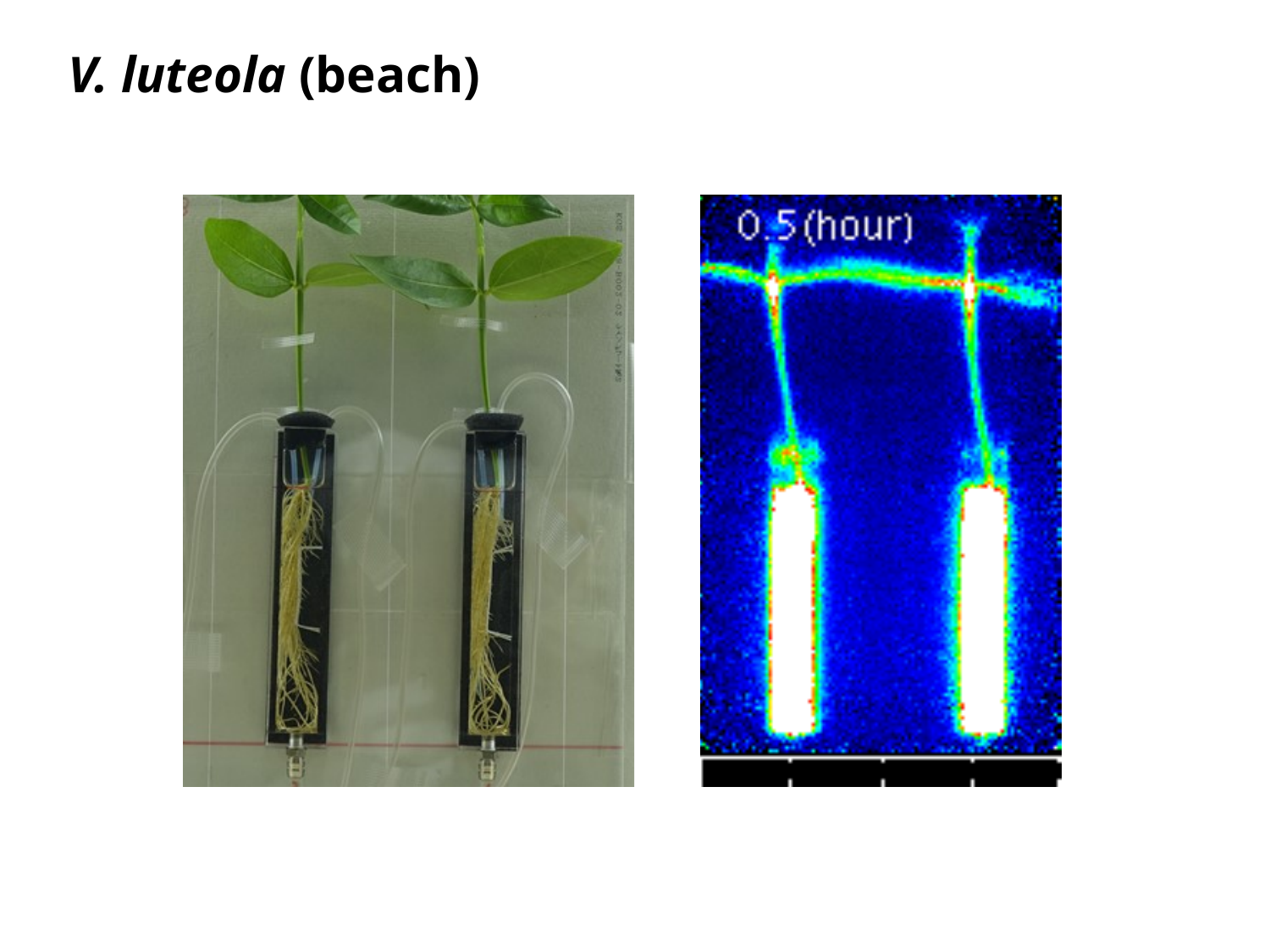

V. luteola (beach)

#### Slide 4
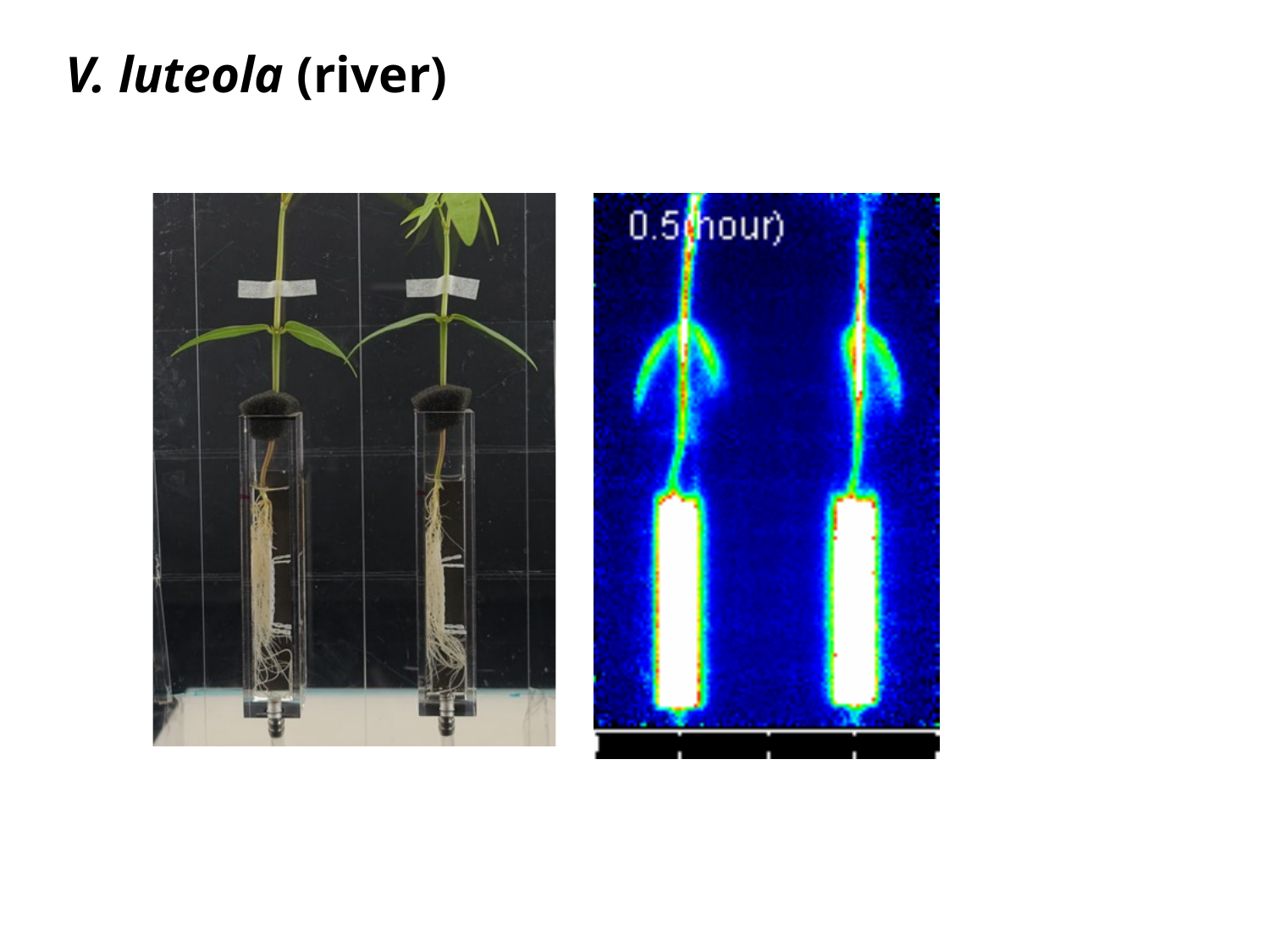

V. luteola (river)

#### Slide 5
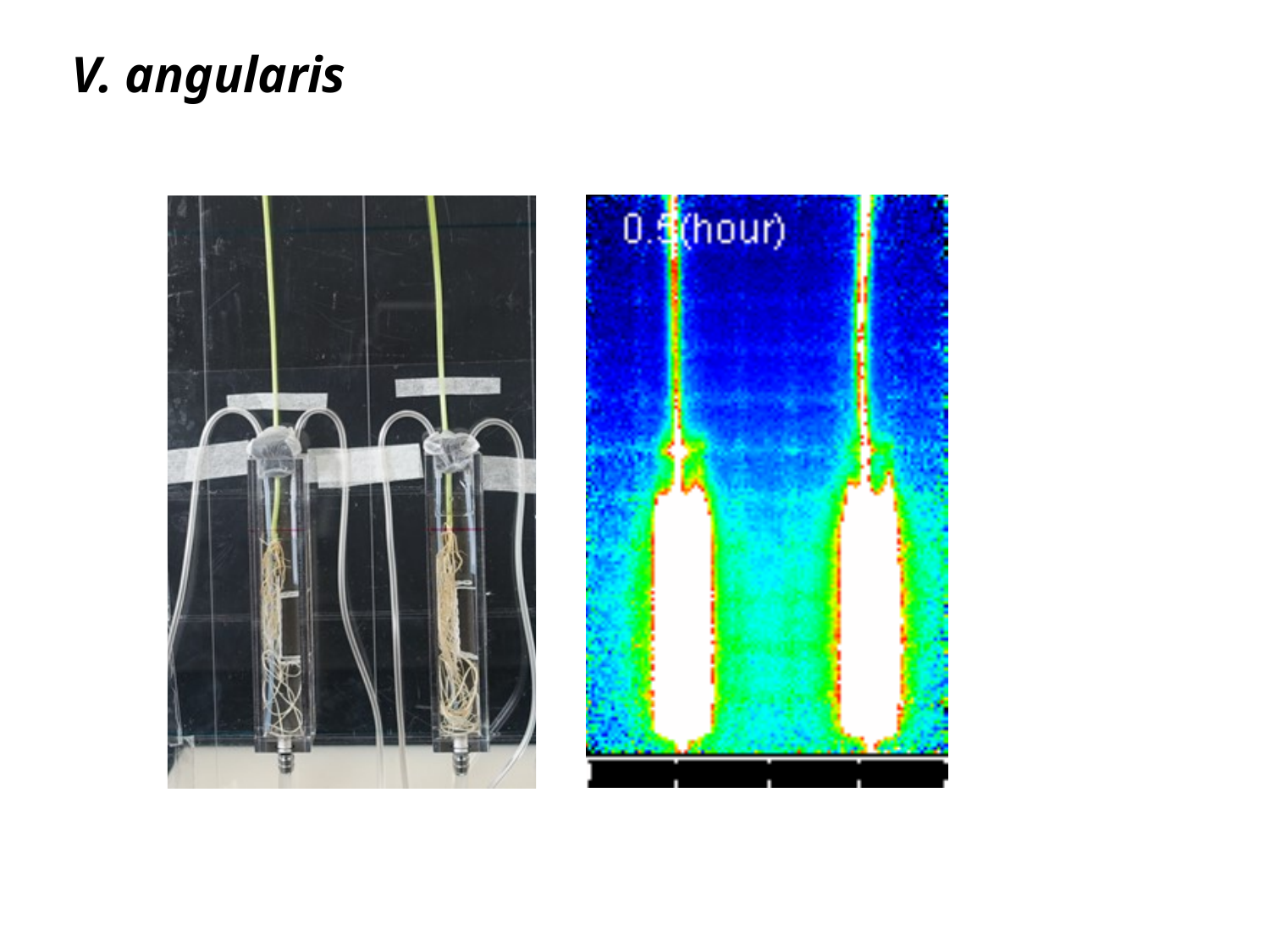

V. angularis
